# Exploring dynamics and network analysis of spike glycoprotein of SARS-COV-2

**DOI:** 10.1101/2020.09.28.317206

**Authors:** Mahdi Ghorbani, Bernard R. Brooks, Jeffery B. Klauda

**Affiliations:** Department of Chemical and Biomolecular Engineering, University of Maryland, College Park, MD 20742, USA; Biophysics Graduate Program, University of Maryland, College Park, MD 20742, USA; Laboratory of Computational Biology, National, Heart, Lung and Blood Institute, National Institutes of Health, Bethesda, Maryland 20824, USA

## Abstract

The ongoing pandemic caused by coronavirus SARS-COV-2 continues to rage with devastating consequences on human health and global economy. The spike glycoprotein on the surface of coronavirus mediates its entry into host cells and is the target of all current antibody design efforts to neutralize the virus. The glycan shield of the spike helps the virus to evade the human immune response by providing a thick sugar-coated barrier against any antibody. To study the dynamic motion of glycans in the spike protein, we performed microsecond-long MD simulation in two different states that correspond to the receptor binding domain in open or closed conformations. Analysis of this microsecond-long simulation revealed a scissoring motion on the N-terminal domain of neighboring monomers in the spike trimer. Role of multiple glycans in shielding of spike protein in different regions were uncovered by a network analysis, where the high betweenness centrality of glycans at the apex revealed their importance and function in the glycan shield. Microdomains of glycans were identified featuring a high degree of intra-communication in these microdomains. An antibody overlap analysis revealed the glycan microdomains as well as individual glycans that inhibit access to the antibody epitopes on the spike protein. Overall, the results of this study provide detailed understanding of the spike glycan shield, which may be utilized for therapeutic efforts against this crisis.

## Introduction

Severe acute respiratory syndrome coronavirus 2 (SARS-COV-2) has rapidly spread worldwide since early 2020 and has been considered one of the most challenging global health crises within the century. SARS-COV-2 has caused more than 20 million cases and more than 1 million deaths worldwide as of September 2020.(1, 2) Drug and vaccine development are underway and multiple vaccines have entered clinical trials and some of them are in the last stages of development.(3-5)

SARS-COV-2 is a lipid-enveloped single stranded RNA virus belonging to the beta-coronavirus family, which also includes MERS, SARS and bat related coronaviruses.(6-9) A major characteristic of all coronaviruses is the spike protein (S), which protrudes outward from the viral membrane and plays a key role in the entry of the pathogen into the host cells by binding to human angiotensin converting enzyme-2 (h-ACE2).(6, 10-12) The structure of each monomer of the trimeric spike protein in SARS-COV-2 (Figure 1) can be divided into two subunits (S1 and S2), which can be cleaved at residues 685-686 (furin cleavage site) by TMPRSS protease after binding to host cell receptor h-ACE2.(12) S1 subunit in S trimer includes a N-terminal domain (NTD) and a receptor binding domain (RBD) that is responsible for binding to h-ACE2.(13) S2 subunit contains fusion peptides (FP), heptad repeats (HR1 and HR2), a transmembrane (TM) and a cytoplasmic domain (CP). The homotrimeric spike protein is highly glycosylated with 22 predicted N-linked glycosylation and 4 O glycosylation sites per monomer, most of which are confirmed by Cryo-EM studies.(14-17) Glycosylation of proteins plays a crucial role in numerous biological process such as protein folding and evasion of immune response.(18)

**Figure 1.**
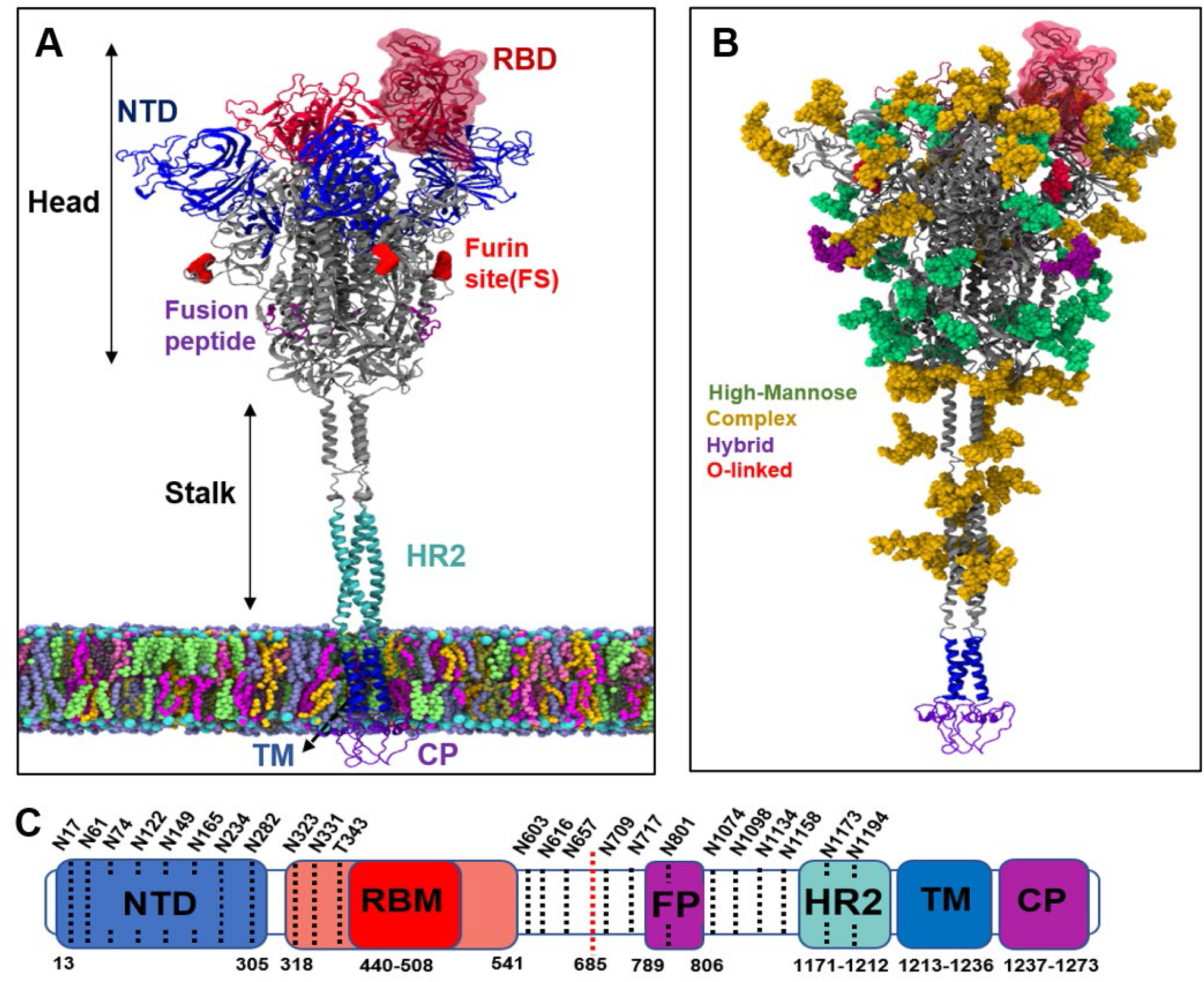
Structure of spike protein and its glycosylation pattern. (a) Different regions of spike protein including N-terminal domain (NTD), receptor binding domain (RBD), Furin cleavage site for cleaving between S1 and S2 subdomains (FS), Fusion peptides (FP), Heptad repeat (HR2), transmembrane (TM) and cytoplasmic (CP) regions. The spike protein is divided into a head and a stalk region. (b) Glycans on the spike protein color-coded based on their types. (c) Sequence of full-length spike protein with domain assignments.

The receptor binding domain (RBD) of SARS-COV-2 binds to its receptor h-ACE2 with a significantly higher affinity than the earlier SARS coronavirus which is suggested to be the reason for its higher infection rate than SARS-COV in 2002-2003.(19) A receptor binding motif (RBM) in RBD makes all the contacts with the receptor by binding its concave surface to the convex exposed surface of h-ACE2.(20) Details of interaction between RBD of SARS-COV-2 and h-ACE2 are studied by us and by other groups, which showed that RBD of SARS-COV-2 binds to h-ACE2 with about 30 kcal/mol higher affinity than SARS-COV RBD. Mutations at critical residues from SARS to SARS-COV-2 RBM are suggested to the be the reason for the higher affinity.(19, 21) Natural mutations at the RBM of SARS-COV-2 have been found in multiple countries, which are not known to be linked with the severity of coronavirus.(22) In an earlier study, we have shown that most of these mutations in RBM do not affect the binding affinity of SARS-COV-2 RBD to h-ACE2 demonstrating the high tolerance of the interface for mutations. Our analysis showed the high contribution of critical residues K417, Q493, Q498 and G502 of RBM in SARS-COV-2 for its high binding affinity. In addition, our alanine scanning of interface residues in nCOV-2019 RBD showed that alanine substitution at some residues such as G502, K417 and L455 can significantly decrease the binding affinity of the complex.(19)

The spike protein in SARS-COV-2 is highly glycosylated with 22 N-linked glycosylation sites and 4 O-linked glycosylation sites.(14, 15) Glycosylation of the spike proteins plays a major role in evading the humoral immune system by cloaking the S protein by a thick shield of glycans.(23, 24) For example, the HIV-1 envelope glycoprotein (Env) features about 93 N-linked glycosylation sites with mostly high mannose glycans (Man5-9), which covers most of the surface of the spike protein in HIV-1 and comprises over half its mass.(25-27) N-linked glycosylation starts with synthesis of precursor oligosaccharides, which are modified to high mannose forms by glucosidases and then trimmed to complex forms in the Golgi by glucosyltransferases for signaling and other glycobiological functions.(28) A higher degree of processing is usually indicative of exposure or accessibility of glycans to enzymes.(25) Dense crowding glycan regions limit the activity of processing enzymes at these locations.

In this study, we report the microsecond long MD simulation of all-atom solvated fully glycosylated spike protein embedded in a viral membrane model in both RBD-up or open state (PDB:6VSB)(10) and RBD-down (PDB:6VXX)(13) or closed state. The structure of the two states for glycosylated S protein in viral membrane were taken from the CHARMM-GUI website for this study.(14) Details of modeling different regions were presented in detail by Im and coworkers.(14) Structural changes in the spike protein happens on the order of microsecond timescale, where a scissoring motion is observed between the NTDs in the RBD-up conformation. We have used network analysis and centrality measures in graph theory to pinpoint the structural features of the glycan shield in the context of glycan-glycan interactions as well as binding of antibodies to the spike protein. A modularity algorithm helped us to find glycan microdomains featuring high glycan-glycan interactions with breaches between microdomains for antibodies to bind and neutralize the virus.

## Results

### Dynamical motions of the spike protein

The root-mean-square deviation (RMSD) for each region of the spike protein in RBD-up and RBD-down states are represented in Figure S1. The stalk region in both RBD-up and RBD-down states show a fluctuating RMSD, which is due to bending motion in this region of spike protein. The tilting in the head of spike is also observed in high resolution cryo-ET images as well as other recent MD studies of glycosylated spike in viral membrane.(29, 30) The bending dynamics in the stalk region is suggested to assist the virus in scanning the cell surface for receptor proteins more efficiently.(30) A snapshot of the open system at 1 µs is represented in Figure 2A, which shows the incline between the head and the stalk domains of spike. The three chains in the spike protein demonstrate various types of motions characterized by different RMSD values (Figure S2). Glycan root-mean-square fluctuation (RMSF) were calculated for heavy atoms in each glycan and then computing the mean and standard deviation for the glycan (Figure 2B for open and Figure S3 for closed state). Consistently, in all chains, few of the NTD glycans such as N74 show high RMSF, which is due to the high solvent exposure of this glycan in NTD compared to other regions. The glycans near the RBD in chain A (RBD-up), such as N234 and T323 showed less fluctuation than other chains. N234 and T323 are sandwiched between RBD and NTD of neighboring monomers in the trimeric spike protein. Glycans in the stalk region showed high RMSF values demonstrating the effective shielding of the spike in this region for both RBD-up and RBD-down states.

**Figure 2.**
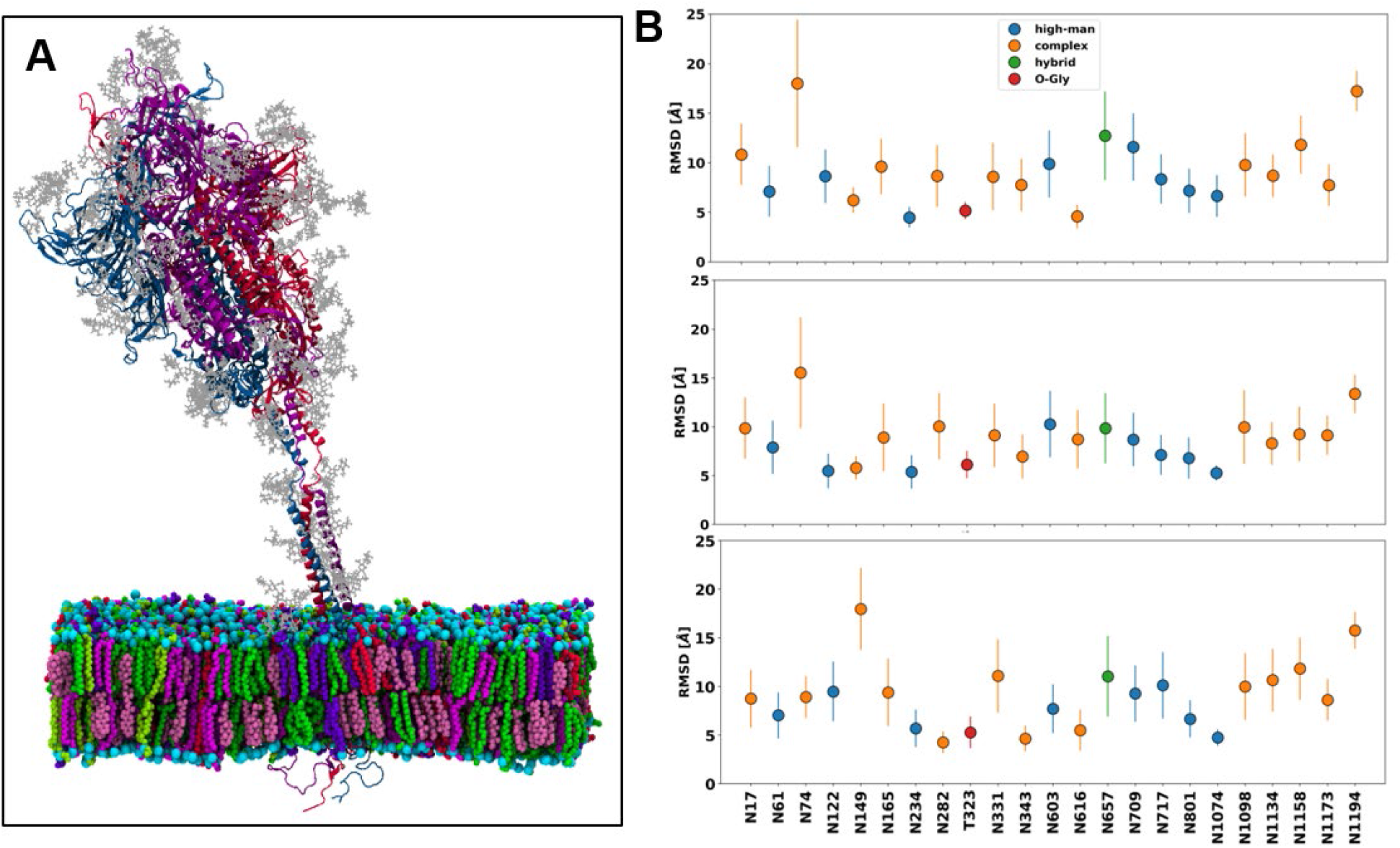
**A)** snapshot of spike in open state after 1000 ns. Different monomers of the spike trimers are color coded with monomer A(up) in red, B in blue and C in purple **B)** RMSF of glycans in the open state of spike for different chains A to C from top to bottom.

A principal component analysis (PCA) was performed on the head region (residues 1-1140) of both RBD-up and RBD-down states to extract the fundamental motions of the trimeric protein. The first two PCs are visualized in 2D plot (Figure 3A and B). Both RBD-down and -up states feature similar behaviors in their PCA plot. In RBD up, PC-1 captures 56% of conformational motion whereas in RBD-down, PC-1 captures only 44% of all conformational motion. This shows the higher conformational change in RBD-up state compared to RBD-down state. The first eigenvector was used to construct the porcupine plots to visualize the most dominant motions in RBD up (Figure S4A) and RBD-down (Figure S4B) states. In the RBD-up state, a scissoring motion is observed between the NTD of chain A and NTD of chain B. Based on the PCA for RBD-up state, the total simulation time was separated into three clusters; cluster-1:0-200ns, cluster-2: 200-600ns and cluster-3: 600-1000ns. The distribution of distance between center of mass of NTD of different chains are calculated for the three different clusters and shown in Figure 3B. In the open state, NTD of chain B goes toward the center of apex. The distribution of distance between the center of apex and each NTD is shown in Figure S4C. In the RBD-down state, PC-1 shows the motion in RBD of chain A toward the open conformation (Figure S4B). The simulation for RBD-down states was separated into two clusters and distribution of the distance between center of mass of RBD and the apex center was calculated for all the chains in both clusters. A clear separation is observed where RBD of chain A is more distant from the apex center in the second cluster (Figure 3D).

**Figure 3.**
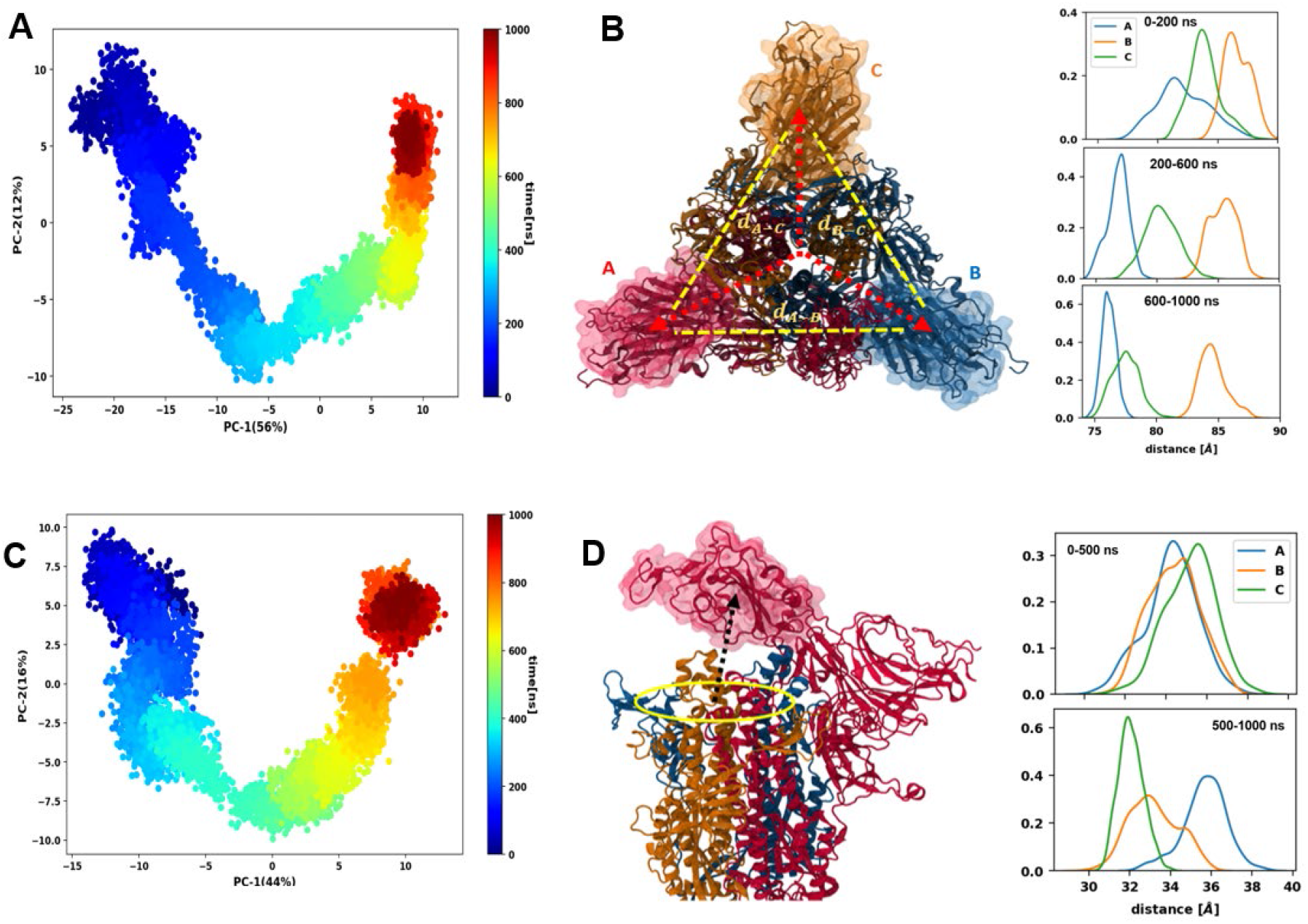
**A)** 2-dimensional (2D) PCA for the open state of spike head **B**) distribution of distance between the center of masses of NTDs on different monomers in the spike trimer for clusters of simulation data 0-200 ns, 200-600 ns and 600-1000 ns. **C**) 2D PCA for the closed spike head **D**) distribution of distance between the center of mass of RBD of different monomer and the center of apex for 0-500 ns and 500-1000 ns of simulation.

### Occupancy of spike protein by glycans

Despite the highly dense glycan shield in the spike protein, there are breaches within the shield that antibodies can bind and neutralize the virus.(31) A volume map of glycans in the spike protein in both up and down conformations are shown in Figure 4, where isosurfaces were visualized for glycans from the 1 µs MD simulation trajectory. The stalk region in the spike protein is highly shielded by the glycans and it is unlikely for antibodies to bind to this region. The spike head on the other hand, shows glycan holes providing opportunities for antibodies to bind. Importantly, NTD of all chains show epitopes free from glycans. The RBM of chain A in the up conformation is completely free from glycans present the least shielded domain of spike. Furthermore, there are regions on the RBD away from the RBM, that show epitopes for antibodies and not shielded by glycans. We have computed the solvent accessible surface area (SASA) of different regions of the spike for antibodies with a probe radius of 7.2 Å that represents the hypervariable domains of antibodies.(31) RBM of chain A in up conformation shows the highest SASA in all chains followed by RBM of chains B and C. On the other hand, RBD of chain A shows the lowest SASA among the three monomers. NTD of all chains show high SASA in all chains. In the closed state, RBM and NTD of chain A shows higher SASA than other chains which is due its conformational change toward the open state. In summary, epitopes on RBM, RBD and NTD of spike protein show high SASA for antibodies to bind and neutralize the spike.

**Figure 4.**
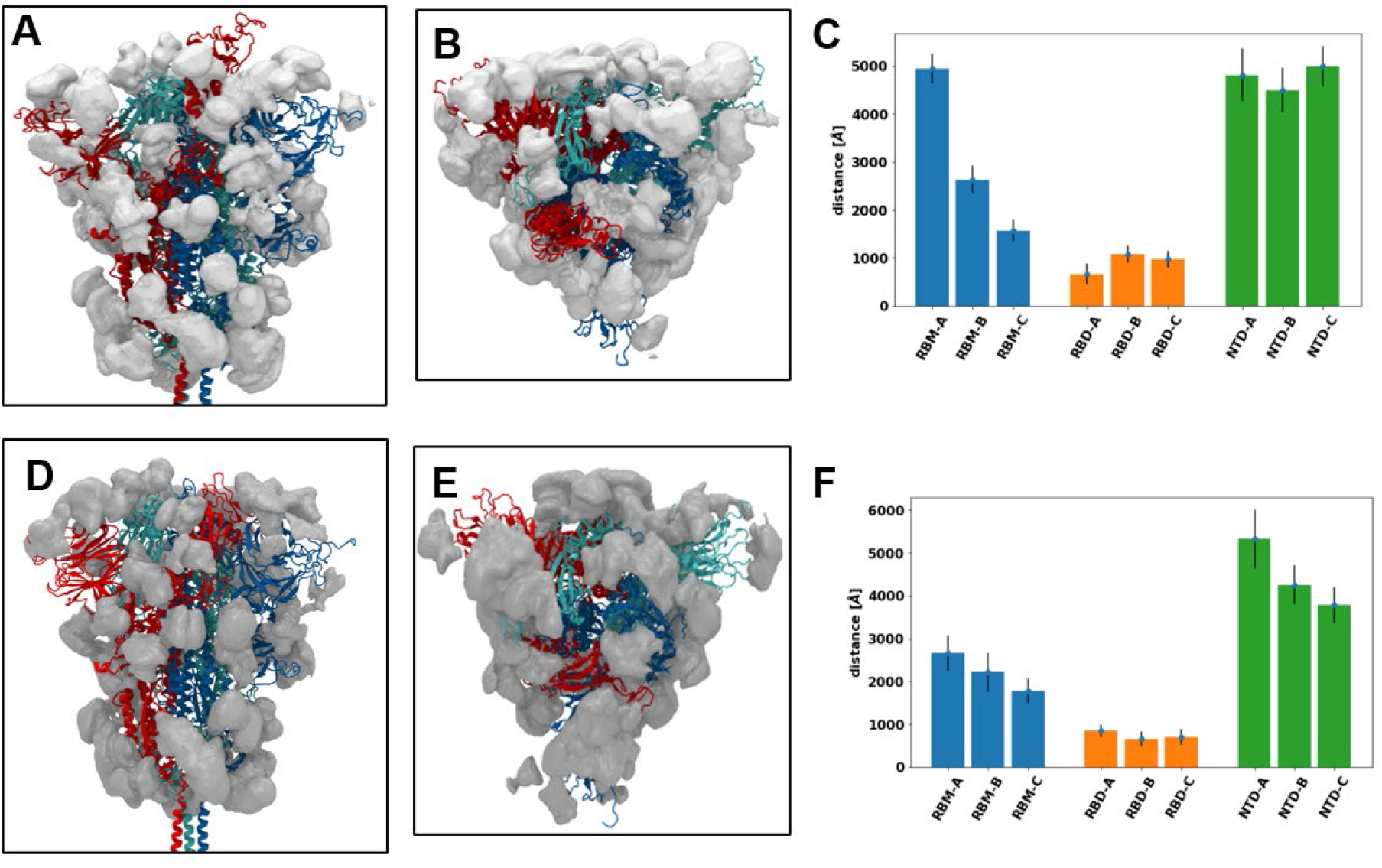
A) glycan occupancy (grey surface) of different regions in the spike head for RBD up state. Monomer A (up) shown as red, monomer B as blue and C as cyan. B) top view of spike head in open state. C) SASA calculated with probe radius of 7.2 Å for different regions of spike head. D) glycan occupancy at closed state. E) top view of closed state F) SASA of closed state.

### Glycan-Glycan interaction and network analysis of glycans

A network analysis was carried out on the glycan shield of the spike protein in both RBD-up and RBD-down states to find the glycans that are most important for an effective shield. This approach has recently been applied to the glycan shield of HIV-1 spike protein.(32, 33) In the glycan network, each glycan represents a node in the graph and two nodes are connected by an edge if the glycans have a distance less than 50 Å in the starting structure. Edges are weighted by the normalized absolute value of non-bonded interaction energy (vdw+electrostatic) between each pair of glycans. A network was built for the whole simulation time in both RBD-up and -down states. All network analysis was performed in networkx and graph visualizations were done using Gephi.(34, 35) The adjacency matrix for these two networks are calculated (Figure S5). Two centrality measurements in graph theory were utilized to analyze the network of glycans. Betweenness centrality quantifies the number of times a node acts as a bridge along the shortest path connecting every two nodes in the graph and is an important indicator of the influence of the node within the network. Eigenvector centrality measures the node’s importance in the network by considering the importance of the neighbors of that node. If the node is connected to many other nodes that are themselves well connected, that node is assigned a high eigenvector centrality score (Figure 5).

**Figure 5.**
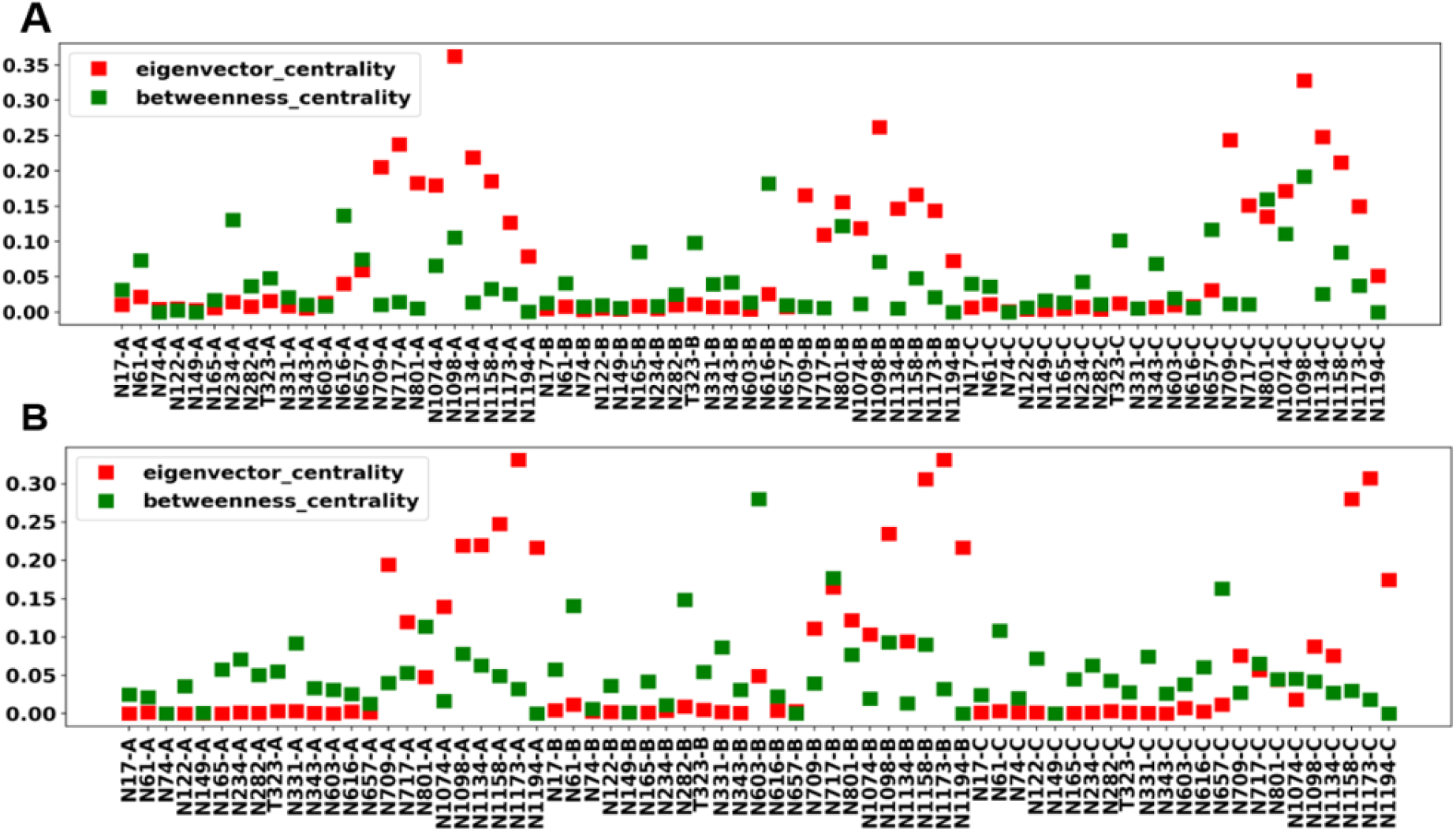
Centrality measurements (eigenvector and betweenness) for A) RBD-up and B) RBD-down states.

Glycans in the stalk region show high eigenvector centrality in both RBD-up and -down states. This means that glycan-glycan interactions in the stalk region is strong and glycans in this region are well connected which result in effective shielding of the stalk against antibodies. In contrast, connections in the spike head and specially at the apex are sparser as the eigenvector centralities are small for this region. Two apex glycans (N234A and N165B) near RBD of chain A (up) in the open state, show a high betweenness centrality (BC). These two glycans show a low BC in the closed state. Glycan N616B in RBD-up and N603B in RBD-down show highest BC in their corresponding network which shows the great impact of this glycan in the proper shielding of the spike protein. Glycans N603 and N616 connect lower head with upper head glycans and are highly central in the network.

Glycans in head region are well separated from glycans that shield the stalk of spike protein. Consequently, we performed network analysis of glycans in the head region of spike protein for the total simulation time. Centrality measurements for open and closed conformations of spike head are shown in Figures S6A and B. A modularity maximization algorithm(36) is used, which resulted in identifying 5 different glycan microdomains for RBD-up (Figure 6A) and 4 glycan microdomains in RBD-down states (Figure 6B). Microdomains feature a high glycan-glycan interaction among them and lower number of edges between different microdomains.(32) The higher number of microdomains in RBD-up state shows that the spike protein is more vulnerable when RBD is in up conformation. Glycans in the lower head all belong to the same microdomain (Cyan-I) in both RBD-up and RBD-down states. This demonstrates the effective shielding of the lower head by glycans regardless of the RBD conformation as all these glycans belong to the same microdomain. Glycan microdomains in RBD-up and -down conformations were mapped onto the spike head to visualize these clusters on the protein (figure 6). Overall connectivity is lower near the RBD as this region is divided into three different microdomains in RBD-down and four microdomains in RBD-up state. In RBD-down state, most glycans have similar BC and only one glycan N603B shows a relatively higher BC in the network. This is a high mannose glycan, which connects the upper head with the lower head region in the glycan network and is crucial for effective shielding. In both RBD-up and -down states, glycans at the lower head also showed high eigenvector centrality, indicating the effective shield of this region. When RBD is in up conformation, glycans near RBD of chain A (up) can interact with glycans from NTD of chain B. As a result, when RBD is open, glycans N234A, T323A, N331A and N343A all belong to the microdomain that comprises glycans of chain B (shown as Green). This leads to encompassing the RBD of chain (A) in RBD-up state by the same microdomain (Green-III), which enhances the shielding of RBD region away from the RBM. This is also demonstrated by the lower SASA of the region of RBD of chain A away from the binding interface with ACE2 (Figure 4B). However, in RBD-down state, the mentioned glycans of chain A are distant from glycans of chain B and therefore they belong to the microdomain that includes other glycans from chain A (shown as Orange-II). Furthermore, glycans N234A and N165B show high BC in RBD-up state, which is due to their interaction with other glycans from the space left open from RBD of chain A. Glycan N616, which is a fucosylated complex glycan, also shows a high BC in RBD-up state. Interestingly, most of the glycans in the lower head region are oligomannose, whereas the glycans at the upper head region (apex) are mostly complex glycans. Complex glycans have a higher degree of processing by glycan processing enzymes. This correlates with the higher number of microdomains in upper head region where connections between glycans are sparser than the lower head region. Glycan sparsity was shown before to correlate with the degree of processing.

**Figure 6.**
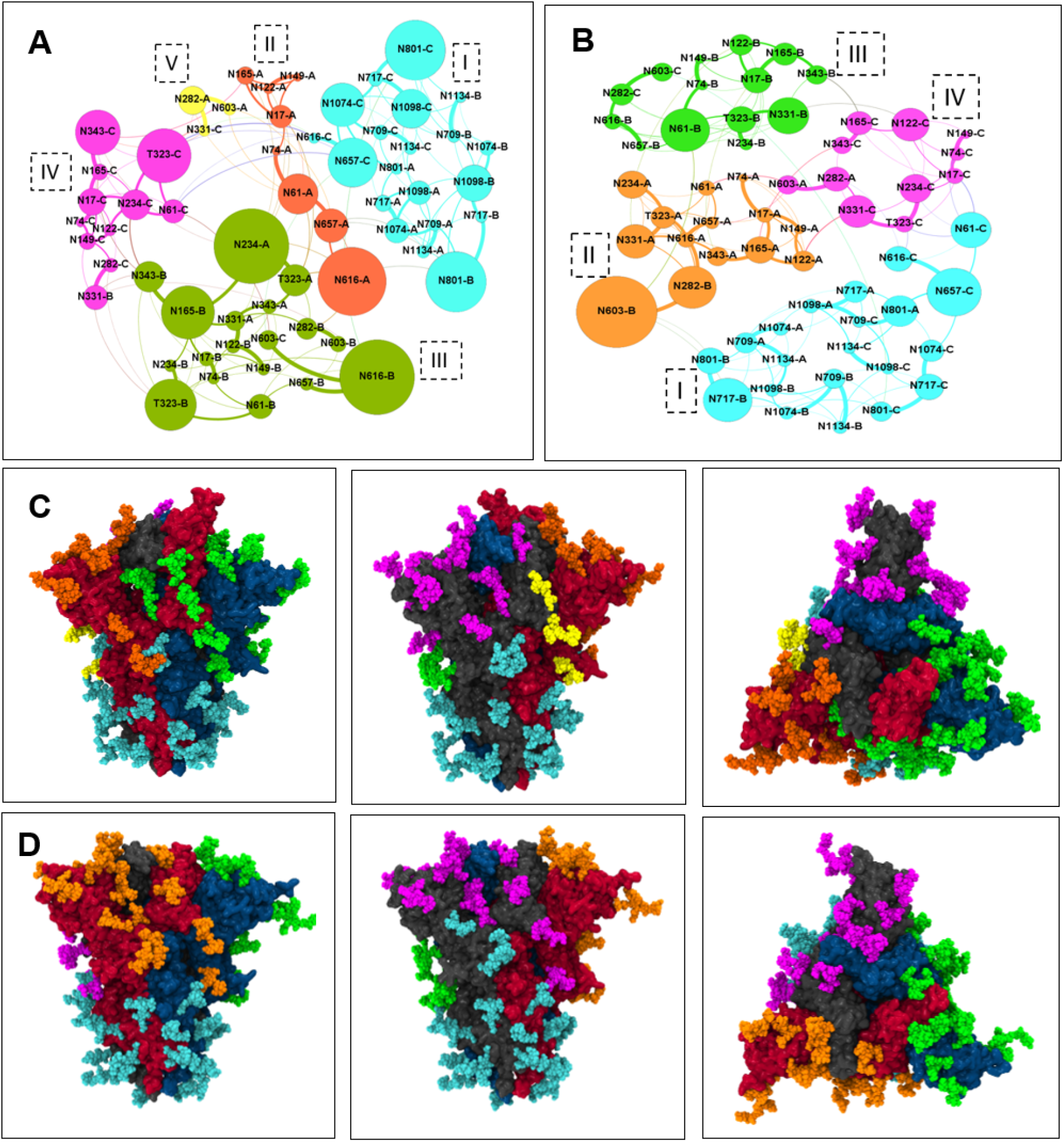
**A**) microdomains in the open state of spike head with each microdomain color-coded. Glycan are connected through and the thickness of the edge shows the edge weight. **B**) microdomains in the closed state of spike head. **C**) microdomains color coded on the spike protein in open state **D**) microdomains in the closed state of spike head.

To identify if the interactions between glycans is coupled with protein conformational changes, we performed the network analysis on the clusters of simulation found from PCA. In the RBD-up state, separate networks were built for 0-200, 200-600 and 600-1000 ns of simulation. Similarly, in RBD-down state networks were built for the first half (0-500 ns) and second half (500-1000 ns) of simulation. BCs were calculated for all the aforementioned networks and changes in betweenness centrality ΔBC were measured between these networks (Figure S6C and D). For the open state simulation, the scissoring motion in NTD is coupled with increasing the BC of N234A, N603A and N165B. As the NTD of chains A and B come closer together glycans in these two chains make stronger connections especially near RBD of chain A where due to the open state, glycan N234A, which is inserted in the space left open by RBD in open conformation, can freely interact with glycans of chain B. N603 is a high mannose glycan in the middle regions of spike and the NTD scissoring motion grants a higher BC for this glycan in RBD-up state. In the RBD-down state, the motion in RBD of chain A to the up conformation increases the BC of glycans such as T323A and N657A and does not affect the BC of most other glycans. Glycan T323A is located at the tail of RBD and N657A is in close proximity of RBD of chain A in the middle head region. The conformational change in RBD brings these glycans closer resulting in a more compact network of glycans in the middle head region and higher BC of these glycans and other neighboring glycans in chain A (Figure S6D).

### Antibody overlap analysis

Neutralizing antibodies look for breaches in the glycan shield, where the glycan densities are sparse.(25, 32) Within a microdomain, glycans are highly connected by a high number of edges in the network and most antibodies bind the regions between these microdomains as comparatively sparse edges connect different microdomains.(25) Therefore, these microdomains help identify susceptible regions of spike protein for immunological studies. Antibodies for spike protein were divided into three categories: RBM-binder, RBD-binder (region of RBD away from RBM) and NTD-binder. The antibodies chosen for each category are presented in the methods section. To investigate the relation between binding of antibodies to known epitopes of the spike protein and the identified glycan microdomains, we utilized an antibody overlap analysis.(25) Antibodies were first overlaid with the spike protein by fitting to their corresponding region in the spike protein. Next, we calculated the average number of clashes between each overlaid antibody and the glycan heavy atoms in each microdomain with a cutoff distance of 5 Å during 1 µs simulation. Results of this analysis are shown in Figure 7A for RBD-up and 7B for RBD-down states.

**Figure 7.**
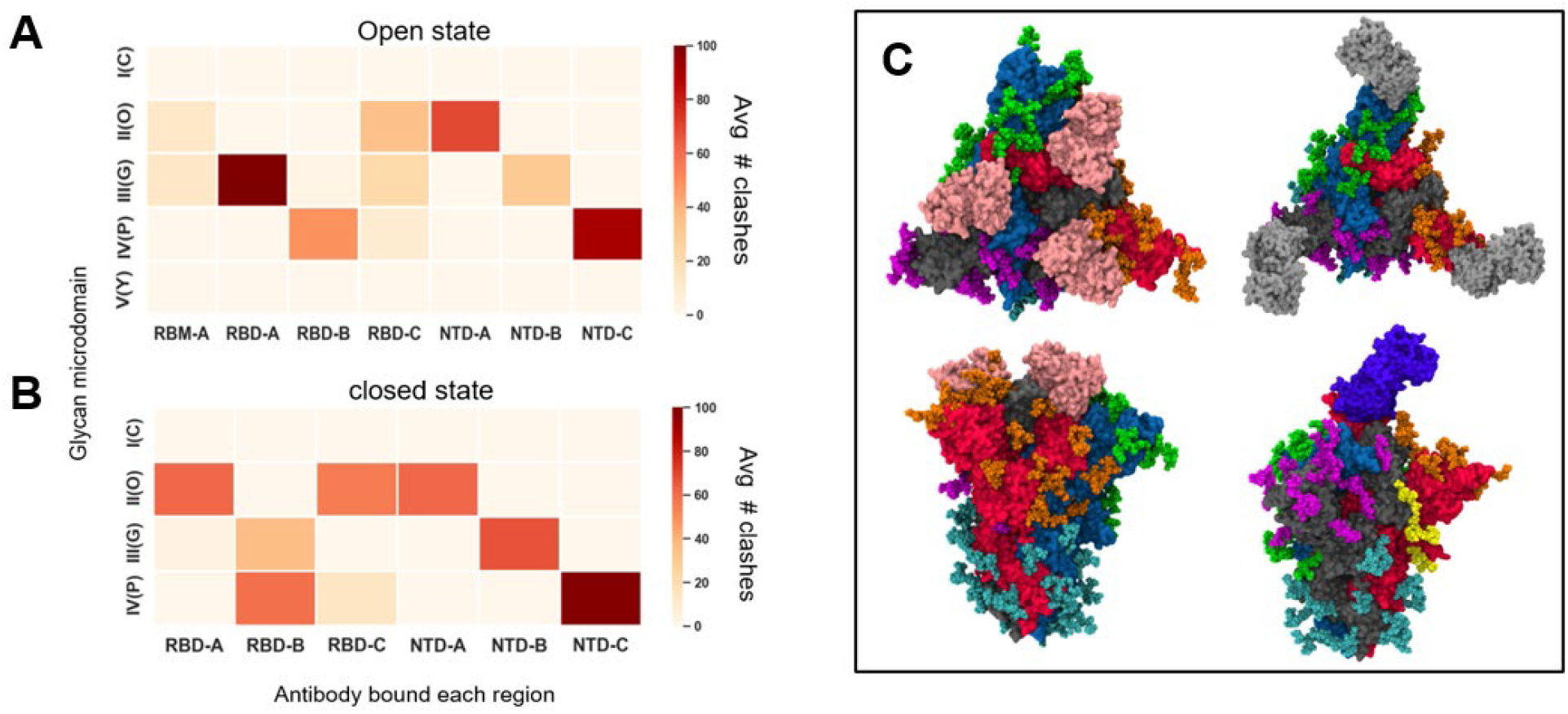
**A)** antibody overlap analysis for the open state. Microdomains are represented by their group and their color in figure 6 with Yellow-V (Y), Purple-IV (P), Green-III (G), Orange-II (O) and Cyan-I (C). The character after each region in x-axis specifies the chain on which the antibody was overlaid, and the number of clashes was calculated **B)** antibody overlap analysis for closed state. Since all the RBD’s are in closed conformation we didn’t calculate clashes for RBM-A **C)** overlaid antibodies with spike protein. Top left RBD binder antibody (pink) with closed state spike. Top right NTD binder antibody(silver) with closed spike. Lower left RBD binder antibody (pink) with open state of spike and lower right RBM binder antibody (blue) with spike open.

The RBM-binder antibody had the lowest number of clashes among all antibodies with microdomains. RBD-binder antibody of chain A had the highest number of clashes among all chains with glycans in microdomain III(G). RBD-binder antibodies of chains B and C have lower number of clashes with glycan microdomains than chain A, where RBD is in the up state. Antibody binding to NTD of chain C (NTD-C) in open state shows a high number of clashes with microdomain IV(P). Similarly, in the closed state, NTD-binder antibody in chain C (NTD-C) also shows a high number of clashes with cluster IV(P). NTD of chain C also showed a lower SASA than NTD of other chains. In the open state, microdomain III(G) comprises the high BC glycans N234A, N165B and T323B. The high BC of glycans in this microdomain correlates with its high number of clashes with antibodies of RBD-A and NTD-A. In RBD-down state, RBD-binder antibodies seem to have similar number of clashes with different glycan microdomains. To identify glycans that have most effect on antibody binding, we also quantified the number of clashes of antibodies with each glycan in different chains and averaged over the simulation time of 1µs (Figure S7). RBM antibody has only a low number of clashes with N165A glycan. In open state, RBD-A antibody has a high number of clashes with multiple glycans (N122B, N149B, N331A and N343A). RBD antibodies bound to the other chains (RBD-B and RBD-C) show lower number of clashes with glycans N122 and N165. NTD antibody of chain C (NTD-C) in both open and closed states show a high number of clashes with glycans N74C and N149C.

## Discussion

Understanding the structure and dynamics of glycan shield in the spike protein of SARS-COV-2 is an indispensable requirement for any antibody and vaccine design endeavors.(23, 37, 38) To this end, we have performed MD simulations of fully glycosylated spike protein of SARS-COV-2 in both open and closed states. Analysis of dynamics for 1 µs trajectories showed a tilting motion in the stalk region, which was also demonstrated with experimental cryo-ET images(29, 30) and suggested to aid the virus with screening the host cells for receptor proteins (Figure 2A). Glycan motions were characterized by RMSF (Figures 2B and S3), which showed higher values for stalk glycans demonstrating the high shielding potential of this normally solvent-exposed region. PCA of the head region of open state of spike demonstrated a scissoring motion between NTDs of neighboring monomers A(up) and B. This scissoring motion resulted in trimer asymmetry with NTD of monomer B advancing toward the center of the apex region and NTD of monomer A showing an angular motion centering on the apex center and toward NTD of monomer B. Based on the PCA, the simulation was divided into three different clusters with distribution of distances between the center of masses of NTDs of different monomers showing different asymmetrical trimers in each cluster (Figures 3B and S4A,C). A scissoring motion in the trimeric spike of HIV-1 on the sub-microsecond timescale was observed by Leminn et. al,(32) which was suggested to be essential for receptor binding. The NTD scissoring motion is only observed in the open state and this could suggest, the scissoring motion is a means employed by the virus to camouflage parts of the spike protein (such as regions of RBD excluding the RBM) when RBD is in the up conformation. The first PC for RBD-down state was visualized (Figure S4C) and shows the conformational change of RBD in chain A from down toward up conformation. Distribution of distance between center of mass of RBD in each monomer from the center of apex region (Figure 3D) exhibited this motion for two clusters of data 0-500ns and 500-1000 ns separated based on the PCA of open state simulation.

The abundance of information in MD simulations of glycosylated spike protein may hinder identifying important biological features of the glycan shield. Therefore, a network analysis approach is used to identify collective behavior of glycans. The most central region of spike based on eigenvector centrality of the network, is shown to be the stalk domain and the lower head region of spike where a dense array of glycans gives rise to resilience to enzymatic actions. Most of the glycans at the lower head region and upper stalk domain are high mannose with the lower degree of processing which correlates with their high centralities in the graph. High eigenvector centrality of lower head and the stalk glycans also makes it hard for neutralizing enzymes to target and to date, no epitopes for antibodies have been found that target this region.(17, 23, 39)

Glycans on the head region demonstrated different behaviors depending on the RBD state (up or down conformation). Interestingly two glycan at the open state, N234A and N165B show a high BC. N234A occupies the volume left open by the RBD in the open state, whereas in the closed state it is directed toward the solvent. The highly conserved glycan N165B is inserted between RBD of chain A(up) and NTD of chain B and in the open state it occupies the volume of left vacant by RBD (A) in the up conformation. Amaro and coworkers(31) studied the fully glycosylated spike protein of SARS-COV-2 computationally and showed that N234A and N165B are crucial for stabilizing the RBD in the up conformation in the open state of spike. Their simulation showed that mutating N234 and N165 to Ala destabilized the RBD in the open state. Furthermore, experimental negative stain electron microscopy and single particle cryo-EM showed that the equilibrium population ratio between the open and closed state is 1:1.(40) Deletion of N234 glycan shifts this ratio to 1:4 favoring the closed state and deletion of glycan at N165 increased the population of open state with a ratio of 2:1. It was shown that N234 glycan stabilizes the open state of the RBD and inhibits the up-to-down conformational change and N165 glycan sterically inhibits the down-to-up conformational change of RBD. Here we have shown that these two glycans in the open state exhibit a high BC, which is due to their interaction with each other as well as other neighboring glycans at RBD of chain A as well as NTD of chain B. We have further demonstrated that the BC of glycans in the head region is coupled with the scissoring motion between NTDs of monomers A and B. In addition, the scissoring motion gives rise to high BC for glycans in the middle region of spike for glycans N616A and N616B. This is caused by the tighter packing of glycans in the asymmetric trimer. In the closed state, the BC of most glycans do not change, which correlates with lower fluctuation of RBD-down simulation.

Modularity maximization in network analysis allowed us to find 5 microdomains of glycans in RBD-up and 4 microdomains in RBD-down states. The higher number of microdomains at the apex of RBD-up state shows the more vulnerability of spike protein to antibodies in the open state. Glycans at the lower head in both open and closed states belong to the same microdomain (Cyan-I), which shows large number of edges between glycans in this region and thereby effective shielding. Apex glycans are divided into three microdomains in closed and four microdomains in open state. The antibody overlap analysis showed that RBM binder antibody in open state (up) shows the lowest number of clashes with glycan microdomains (Figure 7A). The RBM of chain A also showed the highest SASA among other epitopes on spike protein which shows its great potential for antibody design strategies. NTD-binder antibodies can also bind to epitopes on the NTD of spike protein. These antibodies showed high number of clashes with glycans N74 and N149 of the respective monomer that they bind. The glycans on the surface of spike protein exert a collective behavior, which is an important property that needs to be considered in the context of vaccine and antibody design.

## Methods

### Molecular dynamics simulation

Structures for glycosylated spike protein in the viral membrane with RBD-up and RBD-down states were taken from the CHARMM-GUI website (http://www.charmm-gui.org/docs/archive/covid-19).(14) Details of modeling the missing regions in the spike protein are presented by Im et. al. (14) CHARMM36 forcefield(41-43) was used for protein, lipids and carbohydrates in this study. The glycan composition for each site in the spike protein represents the most abundant based on experimental mass spectroscopy data. The selected glycan sequences include 22 N-linked and 1 O-linked for each monomer of trimeric spike and the composition at each site is shown in Table S2.(15, 16) GROMACS(44) software was used for molecular dynamics (MD) simulation. Energy minimization was performed in 5000 steps using the steepest descent algorithm. A LINCS algorithm in all steps constrained the bonds containing hydrogen atoms. Equilibration was performed with a standard 6-step equilibration scripts from CHARMM-GUI with restraints on protein, lipid and glycan atoms.(45) For the first three steps of equilibration, each step included 125 ps using Berendsen thermostat at temperature 310.15 K and a coupling constant of 1 ps.(46) In the last 4 equilibration steps a Berendsen barostat(47) was used to maintain the pressure at 1 bar. For the production step, all restraints were removed, and the system was simulated under a NPT ensemble using the Parrinello-Rahman barostat(48) with a compressibility of 4.5 × 10^−5^ bar^−1^ and a coupling constant of 5 ps. The temperature was maintained at 310 K using a Nosé-Hoover thermostat with a temperature coupling constant of 1 ps.(49) The production run lasted 1µs for each system (RBD-up and RBD-down) with a 2 fs timestep and the particle-mesh Ewald (PME)(50) for long range electrostatic interactions using GROMACS 2018.3 package.(44)

### Solvent accessible surface area (SASA)

SASA was calculated using VMD(51) with a probe radius of 7.2 Å which represents the hypervariable region of antibodies.(31) Multiple regions were chosen for SASA calculation: RBM (residues 440 to 508). RBD (residues of RBD away from RBM 330-440 and 509-520) and NTD (residues 13 to 310)

### Glycan-antibody overlap analysis

Antibodies that bind and neutralize spike protein are divided into three categories(39): antibodies that bind to the exposed part of RBD (RBM-binder), antibodies that bind to epitopes away from the RBM in RBD or RBD-binder and antibodies that bind to epitopes on NTD (NTD- binder). Three different antibodies were used in this study: B38 for RBM binder (PDB:7BZ5)(52), S309 for RBD binder (PDB:6WPT for open and PDB:6WPS for closed state)(53) and 4A8 antibody as NTD-binder (PDB:7C2L)(54).

## Acknowledgement

This work was partially-supported by National Heart, Lung and Blood institute (NHLBI) at the National Institute of Health (NIH) for BRB and MG; in addition it was partially supported by the National Science Foundation [grant number CHE2029900] to JBK. The authors acknowledge the biowulf high performance computation center at NIH for providing the time and resources for this project. The authors would like to dedicate this article to the doctors and nurses who sacrificed their time, health and even their lives to fight COVID-19, particularly those in Iran and the United States. JBK would also like to dedicate this work to family friend Joe Kaplan (Silver Spring, MD) who passed away due to COVID-19 on April 22, 2020.

## Supporting information

Additional information in forms of tables and graphs on glycan types, membrane composition, RMSD and RMSF, other network analysis results and details of antibody overlap analysis are presented in the SI section.

## Notes

### Competing Interest Statement

The authors have declared no competing interest.

