## Supplemental Info for "Exploring dynamics and network analysis of spike glycoprotein of SARS-COV-2"

**This PDF file includes:**

Tables S1 and S2,
Figures S1 to S7
References for SI

**Table S1.** Composition of membrane of the viral membrane taken from reference (1).

| Lipid Type | Outer leaflet | Inner leaflet |
|---|---|---|
| DPPC | 48 | 46 |
| POPC | 72 | 69 |
| DPPE | 144 | 138 |
| POPE | 216 | 207 |
| DPPS | 48 | 46 |
| POPS | 72 | 69 |
| PSM | 240 | 230 |
| CHOL | 360 | 345 |

**Table S2.** Glycan composition at each site in this study. Taken with permission from reference 1.

| Site | Composition and Type | Sequence | Alternative Sequence(s) |
|---|---|---|---|
| N61 N122 N603 N709 N717 N801 N1074 | HexNAc(2)Hex(5) High-Mannose | 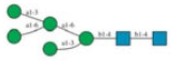 | |
| N234 | HexNAc(2)Hex(8) High-Mannose | 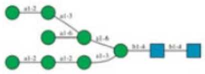 | 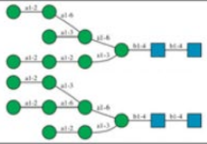 |
| N657 | HexNAc(3)Hex(6) Hybrid | 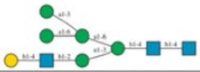 | |
| N149 N331 N343 N616 N1134 | HexNAc(4)Hex(3) Fuc(1) Complex | 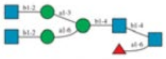 | |
| N1158[a] | HexNAc(4)Hex(4) Complex | 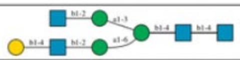 | 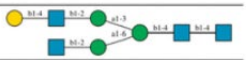 |
| N1098[a] | HexNAc(4)Hex(4) Fuc(1)NeuAc(1) Complex | 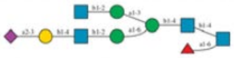 | 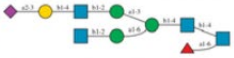 |
| N165[a] | HexNAc(4)Hex(5) Fuc(3)NeuAc(1) Complex | 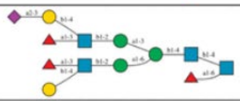 | 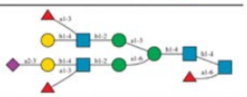 |
| N282 | HexNAc(5)Hex(3) Fuc(1) Complex | 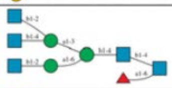 | |
| N17[b] | HexNAc(5)Hex(4) Fuc(1) Complex | 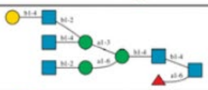 | |
| N1173 | HexNAc(6)Hex(3) Fuc(1) Complex | 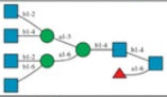 | |
| N74[c] | HexNAc(6)Hex(4) Fuc(1)NeuAc(1) Complex | 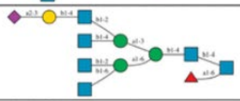 | |
| N1194[d] | HexNAc(6)Hex(5) Fuc(1) NeuAc(1) Complex | 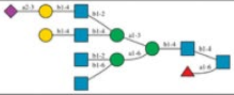 | |
| T323 | O-glycan | 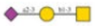 | 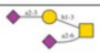 |

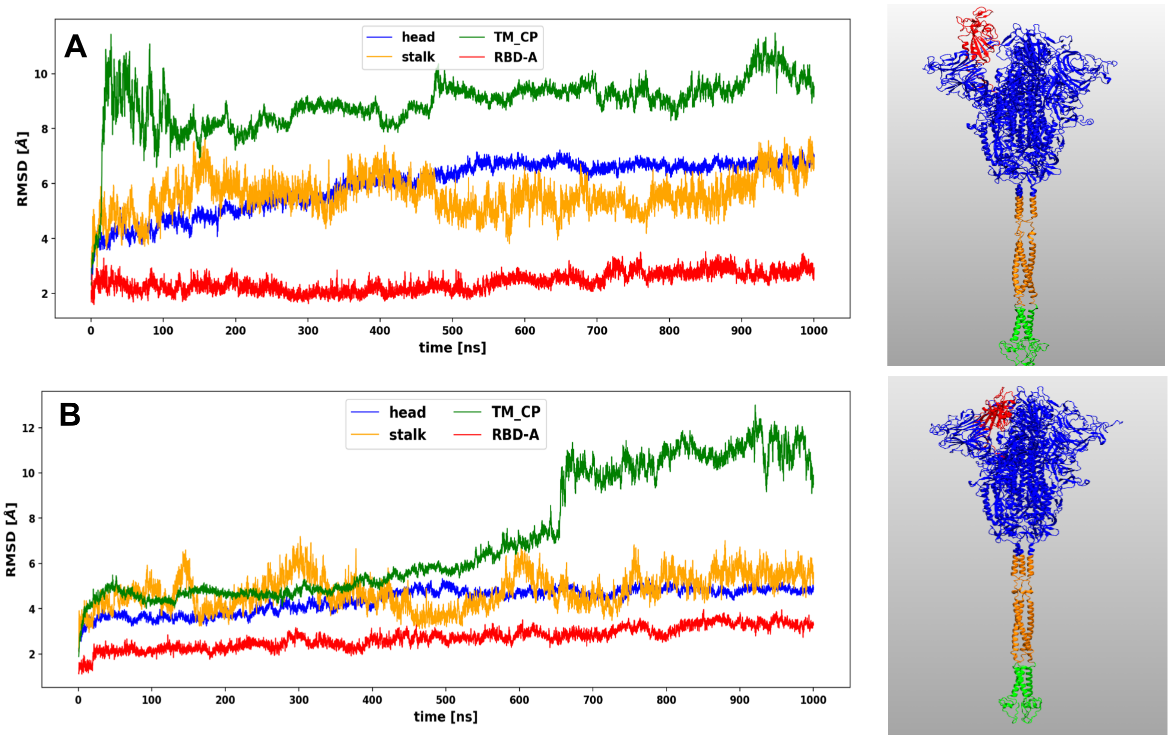


**Figure S1**. RMSD of different regions in the spike protein for A) open and B) closed states

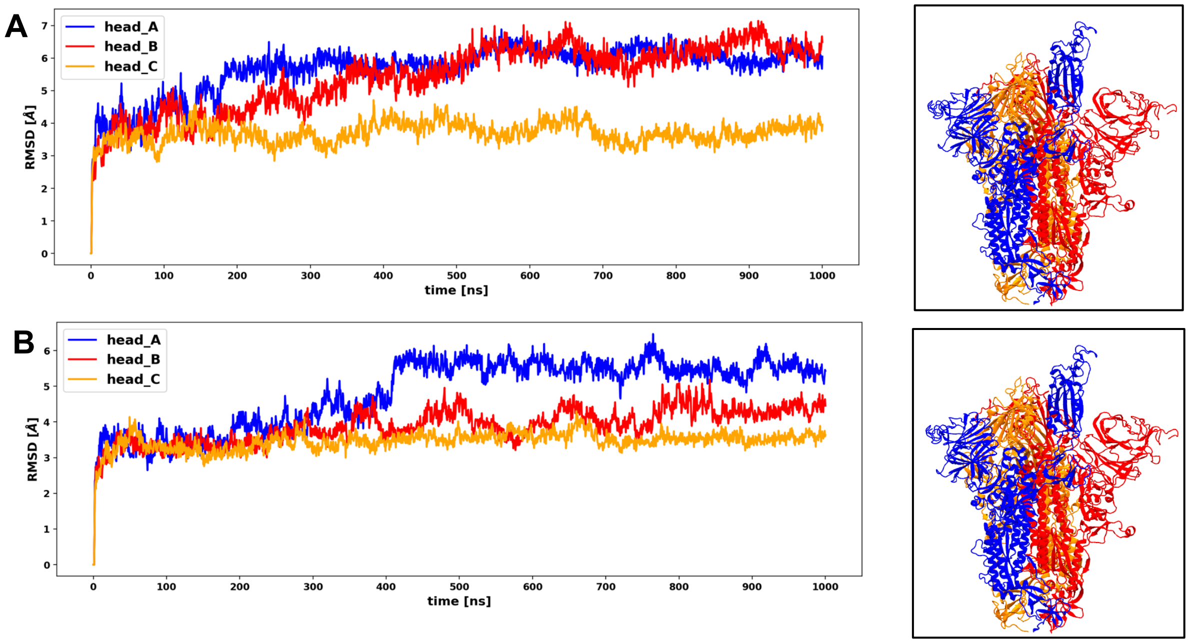


**Figure S2**. RMSD of different monomers in spike trimer for A) open and B) closed conformations

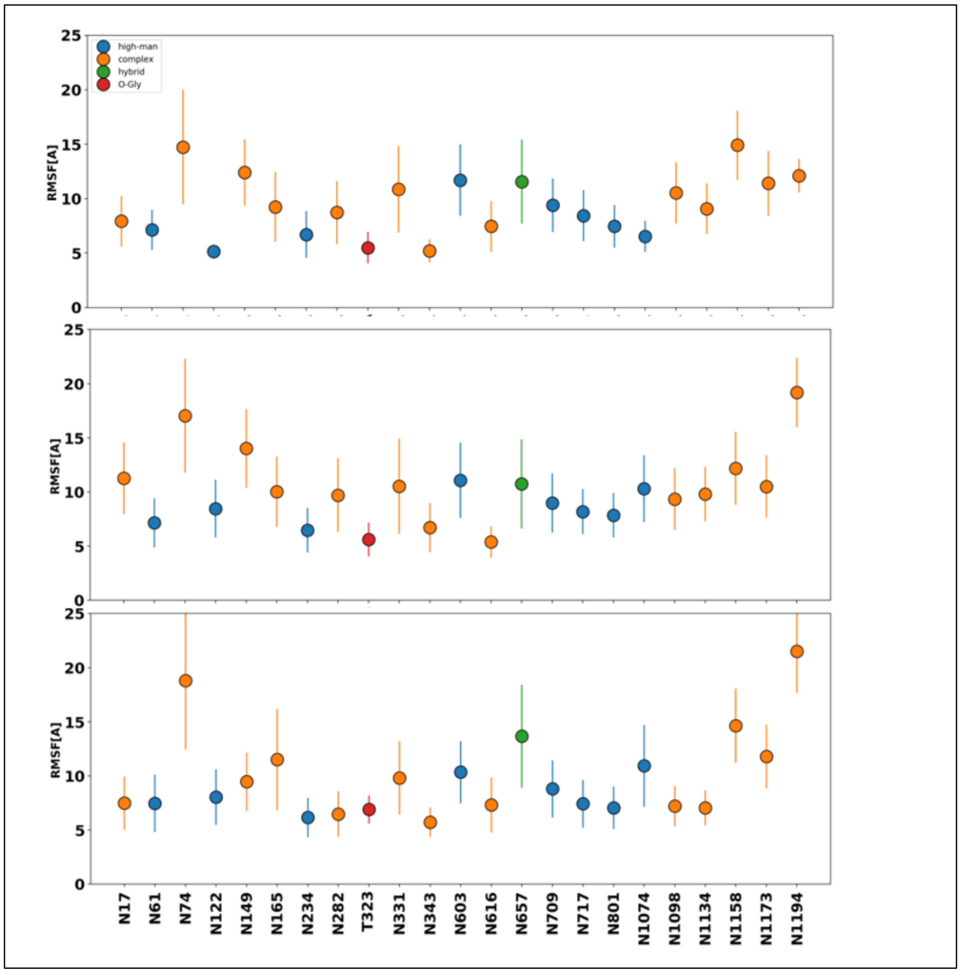


**Figure S3.** Glycan RMSF for closed state and from top to bottom for chain A, B and C.

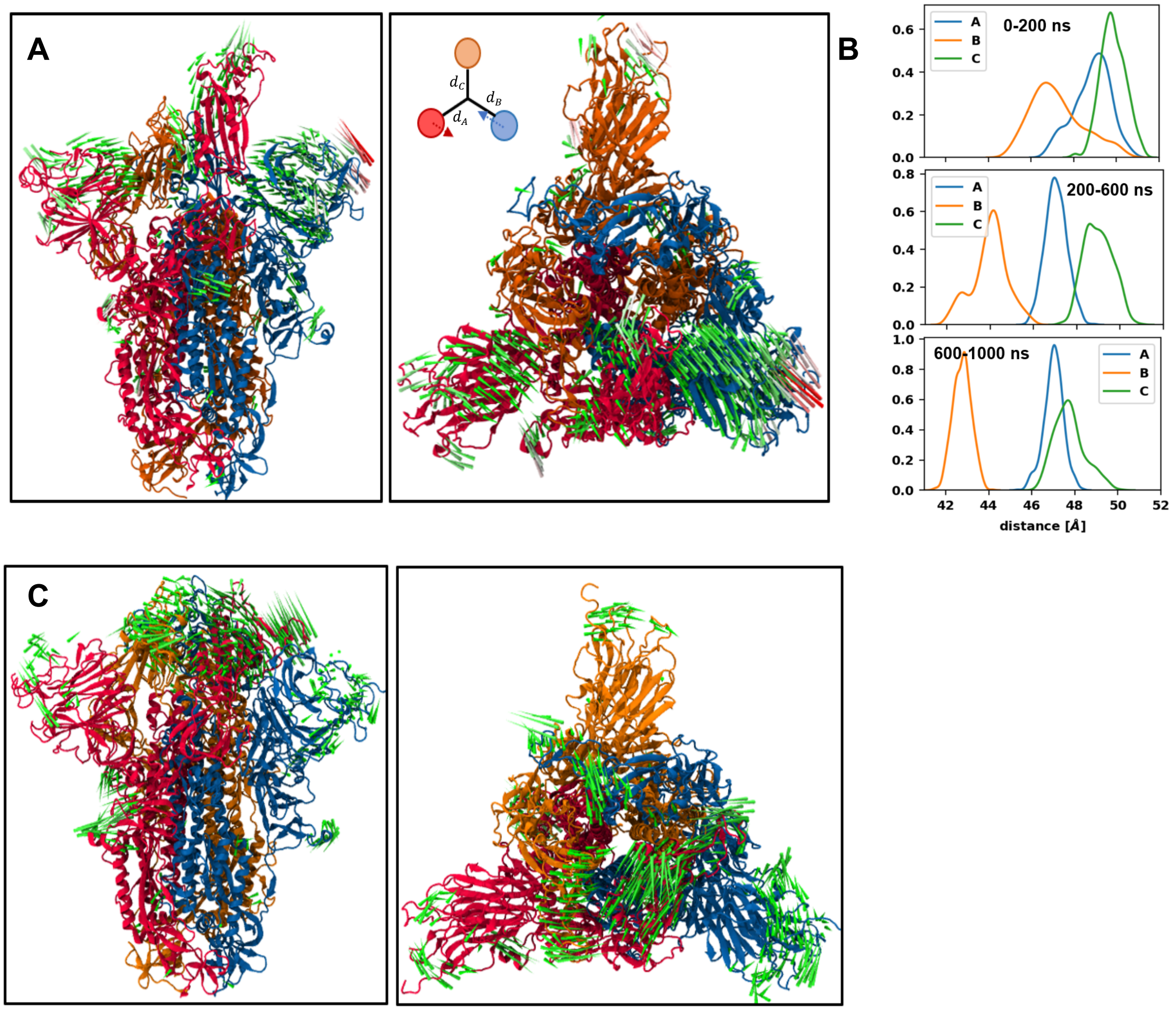


**Figure S4.** A) porcupine plot of first principal component (PC1) for RBD-up state B) distribution of distance between center of mass of NTD of each monomer in open state from the center of apex. From top to bottom corresponds to 0-200 ns, 200-600ns and 600-1000ns of trajectory C) porcupine plot of (PC1) for the closed state.

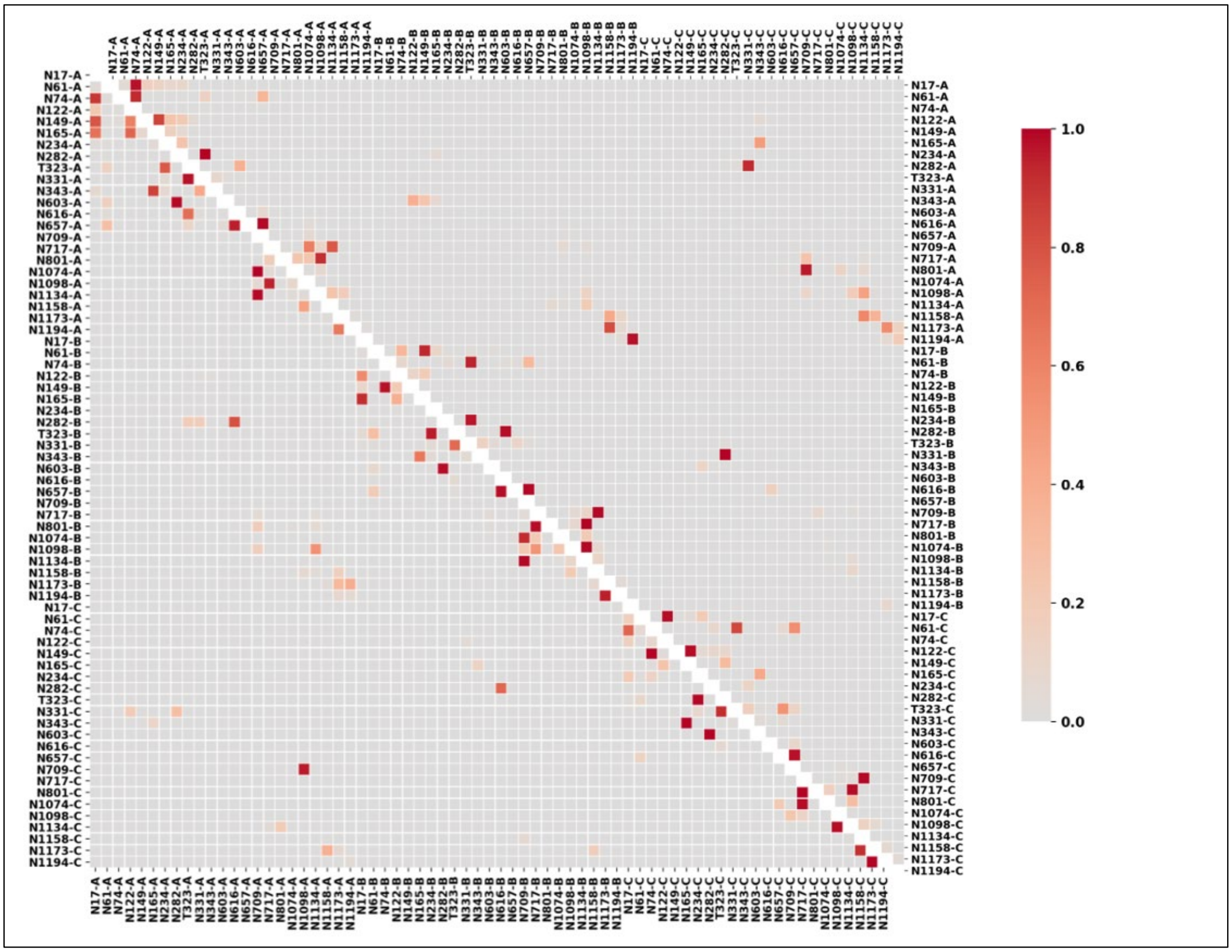


**Figure S5.** Adjacency matrix of glycan-glycan interactions for RBD-up and RBD-down states. Lower triangle represents the adjacency matrix for RBD-down state using the whole simulation and upper triangle is for RBD-up state using the whole simulation.

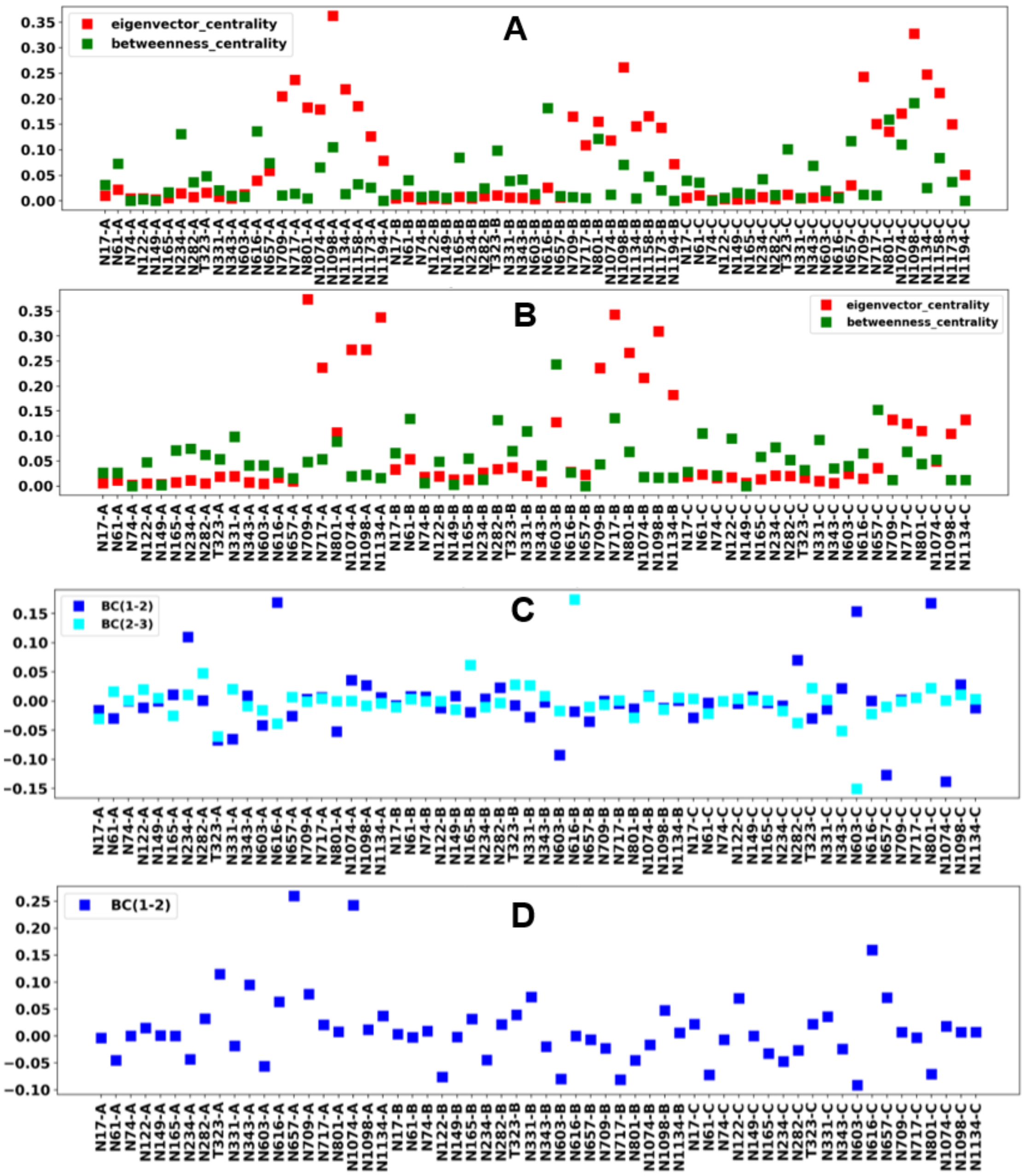


**Figure S6. A)** network centralities for the head region in open state of spike and the total simulation time B) network centralities for the head in closed state and total simulation time. C) change in betweenness centrality of open state between first (0-200 ns) and second (200-600 ns) BC(1-2) and second and third (600 – 1000 ns) BC(2-3). D) change in betweenness centrality of closed state from 0-500 ns to 500-1000 ns.

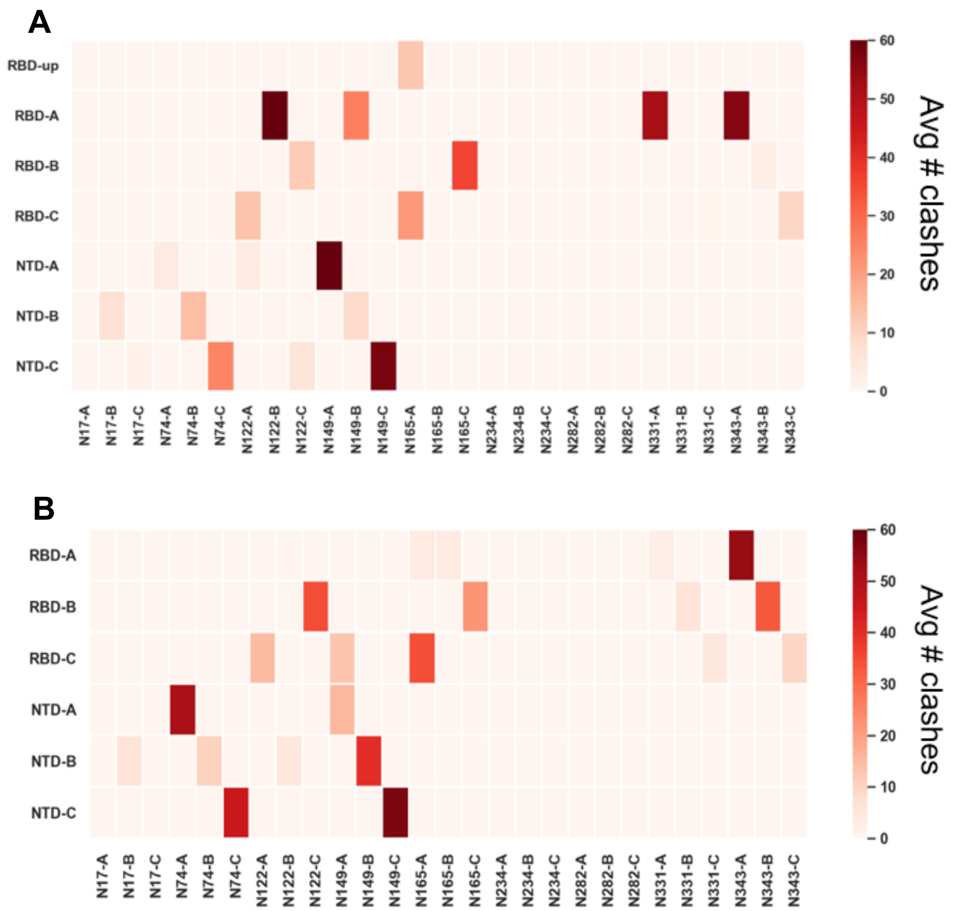


**Figure S7.** Avg number of clashes of each glycan with the overlaid antibody in A) open and B) closed states of the spike
